# Direct long read visualization reveals metabolic interplay between two antimalarial drug targets

**DOI:** 10.1101/2023.02.13.528367

**Authors:** Shiwei Liu, Emily R. Ebel, Aleksander Luniewski, Julia Zulawinska, Mary Lewis Simpson, Jane Kim, Nnenna Ene, Thomas Werner Anthony Braukmann, Molly Congdon, Webster Santos, Ellen Yeh, Jennifer L. Guler

## Abstract

Increases in the copy number of large genomic regions, termed genome amplification, are an important adaptive strategy for malaria parasites. Numerous amplifications across the *Plasmodium falciparum* genome contribute directly to drug resistance or impact the fitness of this protozoan parasite. During the characterization of parasite lines with amplifications of the *dihydroorotate dehydrogenase* (*DHODH*) gene, we detected increased copies of an additional genomic region that encompassed 3 genes (~5 kb) including *GTP cyclohydrolase I* (*GCH1* amplicon). While this gene is reported to increase the fitness of antifolate resistant parasites, *GCH1* amplicons had not previously been implicated in any other antimalarial resistance context. Here, we further explored the association between *GCH1* and *DHODH* copy number. Using long read sequencing and single read visualization, we directly observed a higher number of tandem *GCH1* amplicons in parasites with increased *DHODH* copies (up to 9 amplicons) compared to parental parasites (3 amplicons). While all *GCH1* amplicons shared a consistent structure, expansions arose in 2-unit steps (from 3 to 5 to 7, etc copies). Adaptive evolution of *DHODH* and *GCH1* loci was further bolstered when we evaluated prior selection experiments; *DHODH* amplification was only successful in parasite lines with pre-existing *GCH1* amplicons. These observations, combined with the direct connection between metabolic pathways that contain these enzymes, lead us to propose that the *GCH1* locus is beneficial for the fitness of parasites exposed to *DHODH* inhibitors. This finding highlights the importance of studying variation within individual parasite genomes as well as biochemical connections of drug targets as novel antimalarials move towards clinical approval.

**Author Summary:** Malaria is caused by a protozoan parasite that readily evolves resistance to drugs that are used to treat this deadly disease. Changes that arise in the parasite genome, including extra copies of important genes, directly contribute to this resistance or improve how well the resistant parasite competes. In this study, we identified that extra copies of one gene (*GTP cyclohydrolase* or *GCH1*) were more likely to be found in parasites with extra copies of another gene on a different chromosome (*dihydroorotate dehydrogenase* or *DHODH*). A method that allows us to view long pieces of DNA from individual genomes was especially important for this study; we were able to assess gene number, arrangement, and boundary sequences, which provided clues into how extra copies evolved. Additionally, by analyzing previous experiments, we identified that extra *GCH1* copies improved resistance to drugs that target DHODH. The relationship between these two loci is supported by a direct connection between the folate and pyrimidine biosynthesis pathways that the parasite uses to make DNA. Since *GCH1* amplicons are common in clinical parasites worldwide, this finding highlights the need to study metabolic connections to avoid resistance evolution.

## Introduction

Malaria is a disease caused by the protozoan *Plasmodium* parasite and *P. falciparum* is the leading cause of human malaria deaths (1). Due to the lack of effective vaccines against malaria infection, antimalarial drugs are the primary approach for malaria treatment (2). However, drug efficacy is mitigated by the frequent emergence of antimalarial resistant parasites (3).

Changes in the copy number of large genomic regions, termed copy number variations or CNVs, are an important adaptive strategy for malaria parasites (4–12). Amplification, one type of CNV with increased gene copy number, plays an essential role in the evolution of resistance to various antimalarials (6,7,9,11,13–16). As one example, amplification of the *dihydroorotate dehydrogenase* (*DHODH*) gene directly confers resistance to DHODH inhibitors (i.e. DSM1) in parasites propagated in vitro (11). DHODH is an important enzyme in the *P. falciparum* pyrimidine biosynthesis pathway that contributes resources for nucleic acid synthesis (17,18). *DHODH* amplicons increase transcription and presumably translation of the drug target to directly impact drug sensitivity (11).

In another example, amplification of the *GTP cyclohydrolase 1* (*GCH1*) gene increases the fitness of clinical parasite populations that are resistant to antifolates (i.e. pyrimethamine and sulfadoxine) (4,7,8,16). GCH1 is the first enzyme in the folate biosynthesis pathway and increased flux through this pathway compensates for fitness costs of resistance-conferring mutations in *dihydropteroate synthase* (*DHPS*) and *dihydrofolate synthase* (*DHFS*) genes (4,8,15). Although the contribution of *GCH1* amplicons to antifolate resistance is well studied, this locus has not been reported to contribute to resistance to other antimalarials (15,19,20).

Typically, gene copy number is studied using widely accessible high coverage short read sequencing (11,21–24). However, this approach has limitations including non-unique mapping of repetitive sequences, the inability to resolve complex genomic rearrangements, and the overall difficulty of detecting structural variations (25–27). These challenges are exacerbated by the high AT-content of the *P. falciparum* genome (28,29). Long read technologies such as Oxford Nanopore sequencing have the potential to span low complexity and repetitive regions and more accurately represent structural variation (Cretu Stancu et al., 2017; Sedlazeck et al., 2018a, 2018b; Ho et al., 2020). Moreover, this single molecule sequencing approach allows the examination of clonal heterogeneity within cellular populations (34,35).

In this study, we identified an unprecedented association between *GCH1* and *DHODH* copy numbers from a previously acquired short read data set. To explore this relationship further, we performed long read sequencing and directly observed the step-wise expansion of the *GCH1* amplicon in parasites with elevated *DHODH* copies. Using single long read visualization, we determined that the structure and orientation of the amplicon were preserved during expansion and amplicons increased in a step-wise manner. The positive correlation between *GCH1* and *DHODH* copy number presented here, combined with the recognition that resistance to DHODH inhibitors has only been selected in parasite lines with *GCH1* expansions, suggest the adaptive evolution of these two genomic loci. In addition, the direct connection between folate and pyrimidine biosynthesis supports metabolic interplay between the two pathways. Further study of this relationship is important considering the potential use of DHODH inhibitors to treat clinical malaria with preexisting *GCH1* amplicons.

## Results

Through expanded analysis of short read data from a family of parasites selected with DSM1, originally presented in (11) (**Figure 1A**), we observed a positive association between *GCH1* and *DHODH* copy number (**Figure 1B**). Using droplet digital PCR on the same parasite lines that had recently been propagated in our laboratory, we confirmed that *GCH1* copy number trended higher as *DHODH* copy number increased (**Table 1**).

**Fig 1.**
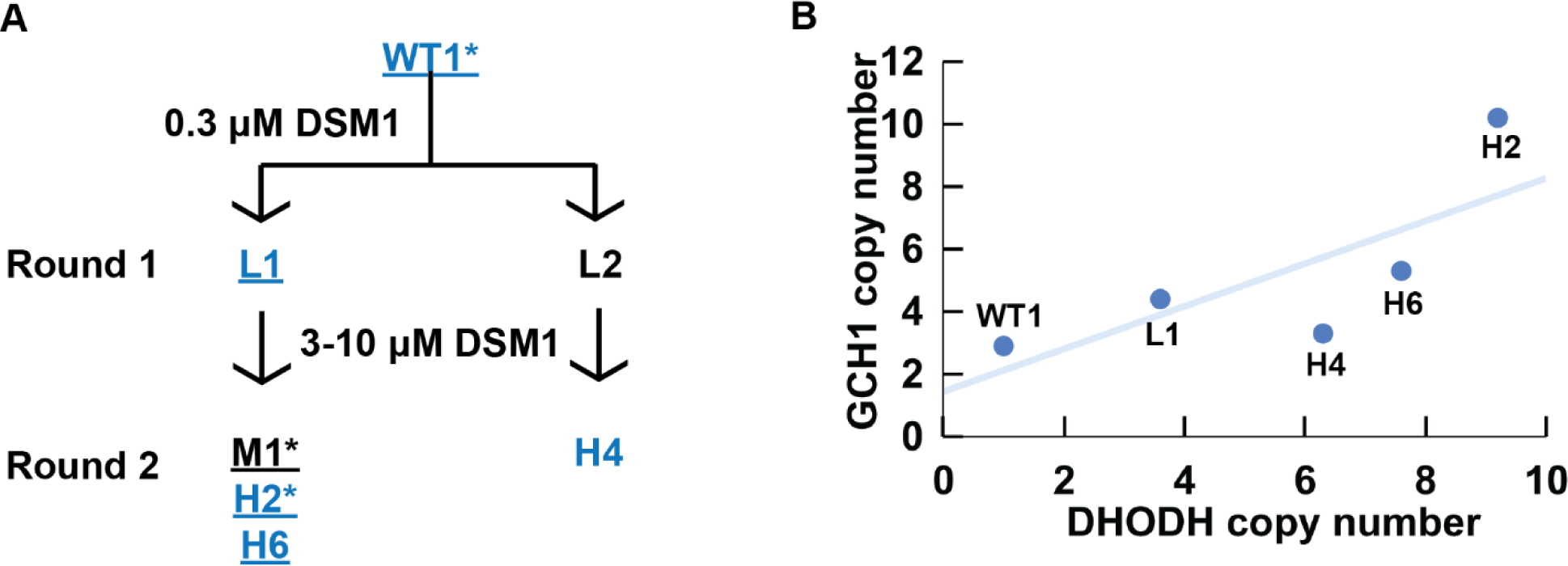
*GCH1* copy number increase is positively correlated with *DHODH* copy number in one family of DSM1 selected parasites. (**A**) DSM1 selection schematic, as presented previously (11). Blue text: Illumina short read sequenced lines. Underline: modern lines confirmed by ddPCR analysis (Table 1). Asterisk (*): lines subjected to long read sequencing in this study. Wild-type (WT1, *Dd2*) *P. falciparum* was selected with DSM1 in two steps; the first step selected for low-level (L) resistance and the second step selected for moderate- (M) or high-level (H) resistance. DSM1 EC50 values are as follows: L1 (1 µM), L2 (0.9 µM), M1 (7.2 µM), H2 (85 µM), H6 (56 µM), H4 (49 µM). All values were previously reported and clone names were adapted as previously (11,36). **(B)** Relationship between *GCH1* and *DHODH* copy number in DSM1 selected parasites as quantified using short read data from Guler et al. 2013. A trend line was added to show the relationship between *GCH1* and *DHODH* copy numbers but a correlation coefficient could not be calculated due to the small sample size (n=5) and dependence among the lines.

**Table 1.** Positive *GCH1*:*DHODH* association is validated using Droplet Digital PCR on modern parasite lines.

| Line | Sample type | Average copy number (SD)* |  |  | *GCH1* CN relative to parent | *DHODH* CN relative to parent |
| --- | --- | --- | --- | --- | --- | --- |
|  |  | *GCH1* | *DHODH* | *ATP6* |  |  |
| WT1 | Parent | 2.5 (0.2) | 0.8 (0.1) | 1.1 (0.0) | 1.0 | 1.0 |
| L1 | Round 1 selection | 3.9 (0.3) | 3.2 (0.1) | 1.0 (0.1) | 1.6 | 4 |
| M1 | Round 1 selection | 3.9 (0.1) | 5.2 (0.3) | 1.0 (0.1) | 1.6 | 6.5 |
| H2 | Round 2 selection | 4.6 (0.2) | 6.0 (0.3) | 1.1 (0.1) | 1.8 | 7.5 |
| H6 | Round 2 selection | 4.3 (0.3) | 7.0 (0.4) | 1.0 (0.1) | 1.7 | 8.8 |

\*The average copy number is calculated by comparing the ddPCR signal to a single copy gene signal (*Seryl tRNA synthetase*, PF3D7\_0717700). N=4. *ATP6* copy number is a single copy gene expected to be 1 in all parasite lines.

We conducted long read sequencing to more precisely define *GCH1* copy number and amplicon structure in the parental line versus two DSM1-selected lines (WT1, M1, and H2, **Table S1**). We directly visualized single reads using an application that represents gene segments of individual long reads (*Materials and Methods*). Small amplicons, like those including *GCH1*, are especially conducive to this approach because long reads span multiple copies of the amplicon as well as both up and downstream regions. Read visualization showed that the general structure of two 3-gene amplicons, separated by an inversion of the same 3-genes, was conserved between the parental and selected lines (**Figure 2A-C**). This amplicon structure was reported previously in the WT1 (*Dd2*) parental line (**Figure 2D**) (4).

**Fig 2.**
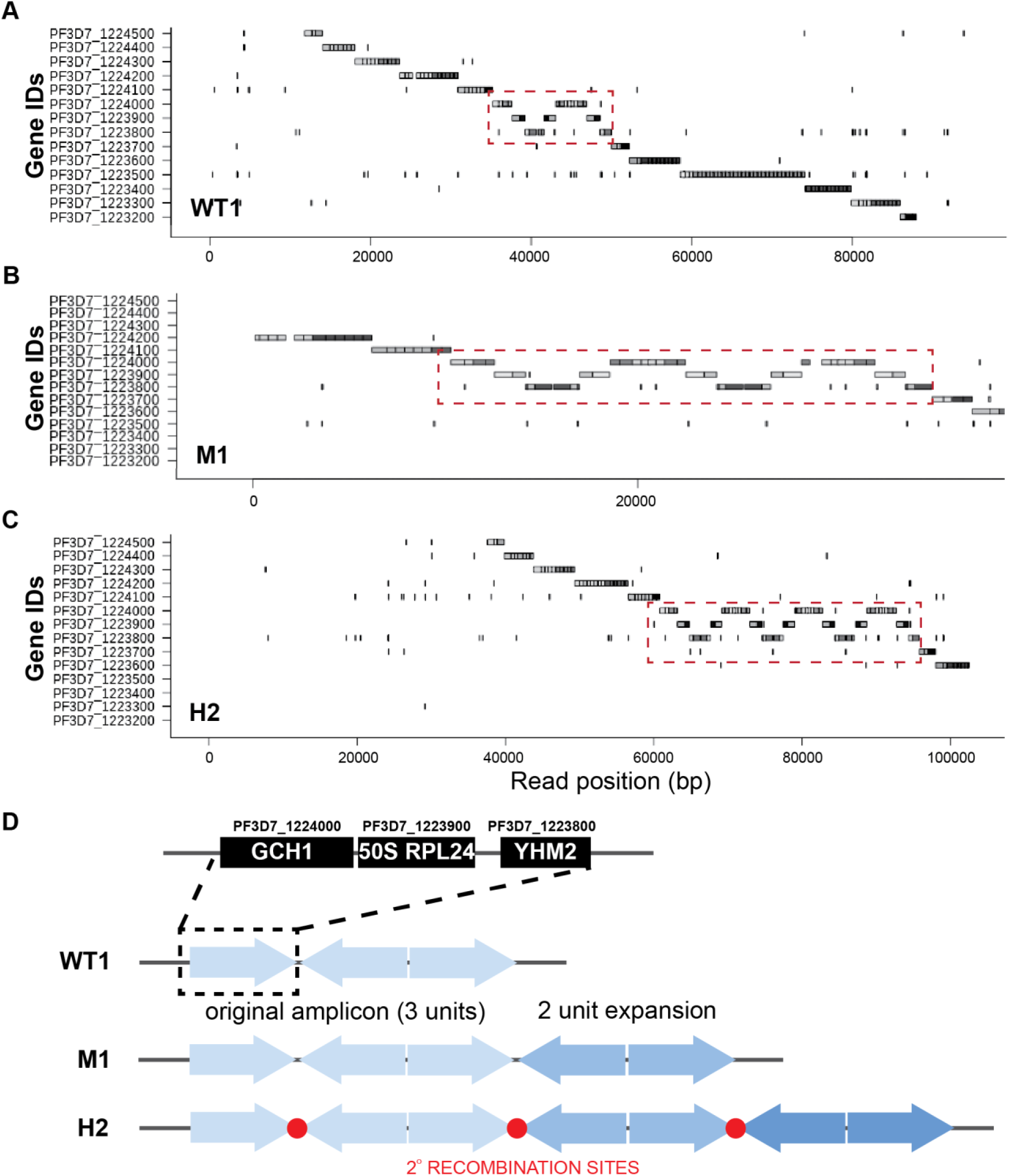
Long-read visualization shows conserved boundaries and structure of *GCH1* amplicon from parental and resistance parasites. **(A-C)** Representative images from the Shiny application comparing the *GCH1* amplicon in WT1 (**A**), M1 (**B**), and H2 (**C**) spanning reads. Red dashed square: copies of *GCH1* amplicon covering 3 genes. Each gene sequence from the *3D7* reference genome (no *GCH1* amplicon represented) was split into <00 bp fragments and blasted against individual Nanopore reads (dark gray: genic regions, light grey: intergenic regions, more details in *Materials and Methods*). **(D)** Orientation of *GCH1* amplicons from each parasite line. The three genes within the *GCH1* amplicon unit include PF3D7_1224000 (*GTP cyclohydrolase I*, *GCH1*), PF3D7_1223900 (*50S ribosomal protein L24*, putative, *50S RPL24)*, and PF3D7_1223800 (*citrate/oxoglutarate carrier protein*, putative, *YHM2*). Note: The gene order (*GCH1-50S RPL24-YHM2*) is reversed compared to the 3D7 reference genome (*YHM2-50S RPL24-GCH1*) in order to facilitate comparison with the read images in panels A-C. Red circles: amplicon junction sequences that act as potential secondary recombination sites.

Using direct read visualization, we also recorded the number of *GCH1* amplicon units (**Figure 2D**) from both spanning and non-spanning reads (**Figure 3A, Table S2**) Spanning reads cover genes both upstream and downstream of the amplicon while non-spanning reads end within the amplicon. We consistently detected up to 3 amplicon units on reads from the parental line. Although the overall mean copy number was similar for all parasite lines (**Figure 3A**, WT1, 2.5 copies; M1, 3.6 copies; H2; 3.2 copies, F(2,95)=[2.61], p = 0.07), we observed more reads that encompassed a higher number of *GCH1* amplicon units in the selected lines (up to 9 units, **Figure 3A**). When we restricted our analysis to spanning reads, or those that displayed the entire amplified region, we detected a significant difference in mean copy number between the parasite lines (WT1 vs M1, p=0.0003 and WT1 vs H2, p=0.01; **Figure S1**). The presence of longer reads was not contributing to this observation; there was not a significant difference in the length of reads used for *GCH1* analysis from each parasite line (F(2,96)=[2.51], p = 0.09, **Figure 3B**, median read lengths: WT1, 51144bp; M1, 49018bp; H2, 26429bp). The larger variation observed for the M1 sample (**Figure 3B**) was due to an improvement in ONT technology and methods leading to an increase in the number of longer reads (i.e. WT1 N50 of 15.6kb on R9.4.1 flow cell and M1 N50 of 99.7kb on R10.4.1 flow cell, **Table S1,** see *Materials and Methods*).

**Fig 3.**
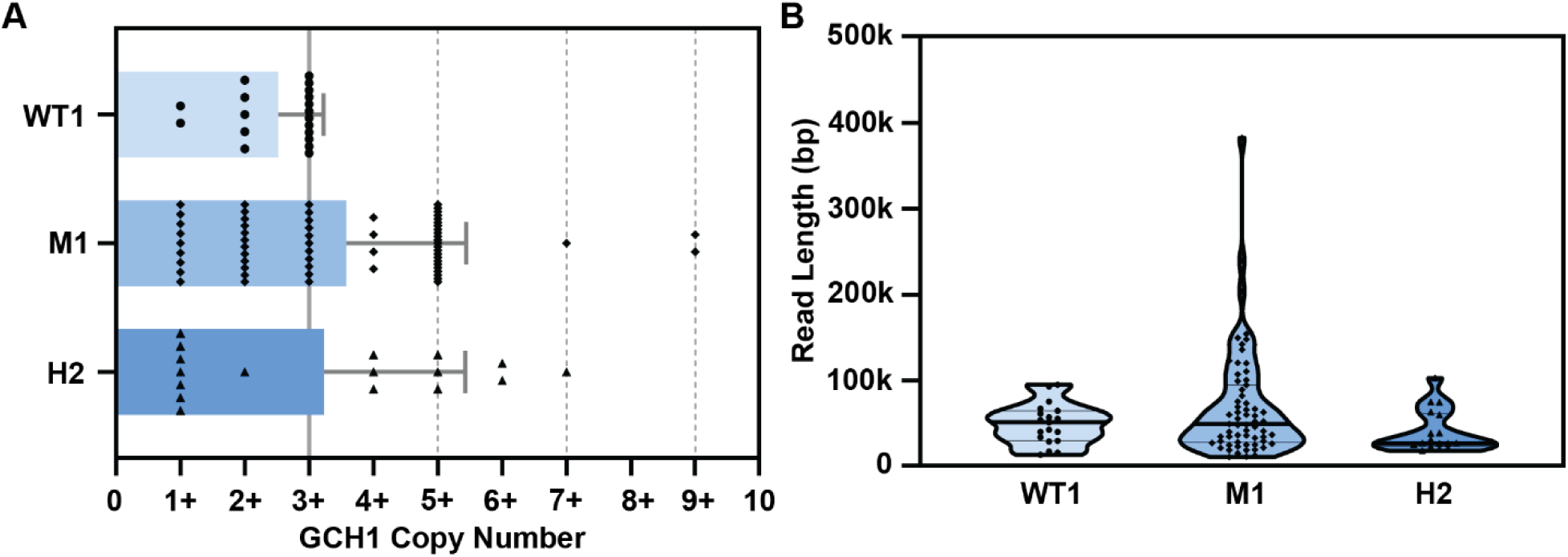
Quantification of long reads displays an increase in *GCH1* amplicon number in DSM1 selected parasites that is not dependent on overall read length. (**A**) *GCH1* copy number from WT1 and selected (M1 and H2) parasite lines. Spanning and non-spanning reads are included; spanning reads were grouped with their corresponding non-spanning count (e.g. a read showing 2 copies of the *GCH1* amplicon is counted as a 2+ read). Bars represent mean, error bars represent standard deviation (dotted lines, values that represent detected spanning reads, see **Figure S1**). (**B**) Read length distribution from long reads (>=10kb) covering the *GCH1* amplicon. Thick line represents median, thin line represents quartiles.

During this analysis, we identified that >50% of reads from M1 and H2 lines depicted amplicon copy numbers greater than the expected WT copy number (**Figure 3A**, **Table S2**). To assess whether variation in the copy number of the *GCH1* locus was common in other laboratory-adapted parasite lines, we evaluated several additional long read datasets (**Table S2**, *Supporting Information,* Vembar et al., 2016). In general, parasite lines exhibited the expected *GCH1* amplicon size (i.e. *Dd2* versus *3D7*, as reported previously (4) and visualized in **Figures 2D** and **S2**) and the amplicon copy number for each line was relatively stable across diverse datasets (limited to <25% of reads above the expected copy number).

When we evaluated the amplicon structure, all M1 and H2 reads covering the *GCH1* region began with a set of 3 amplicon units, as observed in the parental line (**Figure 2A**). This basic structure was followed by groups of 2 units, where one copy was inverted and the other was in normal orientation (**Figure 2D**). This suggests that increases in copy number were arising as two unit steps. We identified further evidence of the two unit amplicon expansion when we limited our analysis to only spanning reads, which allow direct quantification of amplicon copy number. Here, we did not detect any spanning reads that carried *even GCH1* copy numbers (only 3, 5, 7, or 9 copies, **Figure S1**).

To investigate whether an increase in *GCH1* amplicons was contributing generally to resistance to DHODH inhibitors, we evaluated the copy number of *GCH1* amplicons using short read data collected from parasites selected with other DSM compounds in two studies (18,37). Contrary to DSM1-selected parasites (**Figure 1B**), we did not detect increases in *GCH1* amplicon number in parasites selected with DSM265, DSM267, or DSM705 compared to parental *Dd2* or *3D7* parasite lines (**Figure S3**, **Table S3**). For this analysis, we only included parasite lines that acquired *DHODH* amplicons to confer resistance.

The absence of a correlation between *DHODH* and *GCH1* copy number in these other datasets suggests that *GCH1* may not play a direct role in resistance. However, parental lines used in these two studies harbored pre-existing *GCH1* amplicons (~3-6 copies, **Figure 4A**), indicating that this locus may support the evolution of resistance involving *DHODH* amplification. Therefore, we compiled information on all previous resistance selections conducted in parasite lines that naturally vary in *GCH1* copy number (7 total studies, **Table 2**). *3D7* and *Dd2* parasite lines, which carry the highest *GCH1* copy numbers (3-6 copies) with the smallest overall amplicon sizes of common laboratory parasites (1 and 3 genes, respectively, **Figure 4B**), were most commonly used in successful resistance selections (6 and 4 studies, respectively, **Table 2**) and were capable of acquiring *DHODH* amplicons (**Figure 4A**). *K1* parasites harbor a larger low copy *GCH1* amplicon (anticipated as 2 copies of 7 genes, **Figure 4B**) and although resistance could be selected in one study, it was 1000-fold less efficient than *Dd2* selections run in parallel (38). The *DHODH* copy number status of this K1-derived selected line was not investigated. Finally, *Hb3*, which harbors one *GCH1* copy (**Table 2**, **Figure 4B**), was incapable of developing resistance in two independent studies (11,38) (**Figure 4A**).

**Fig 4.**
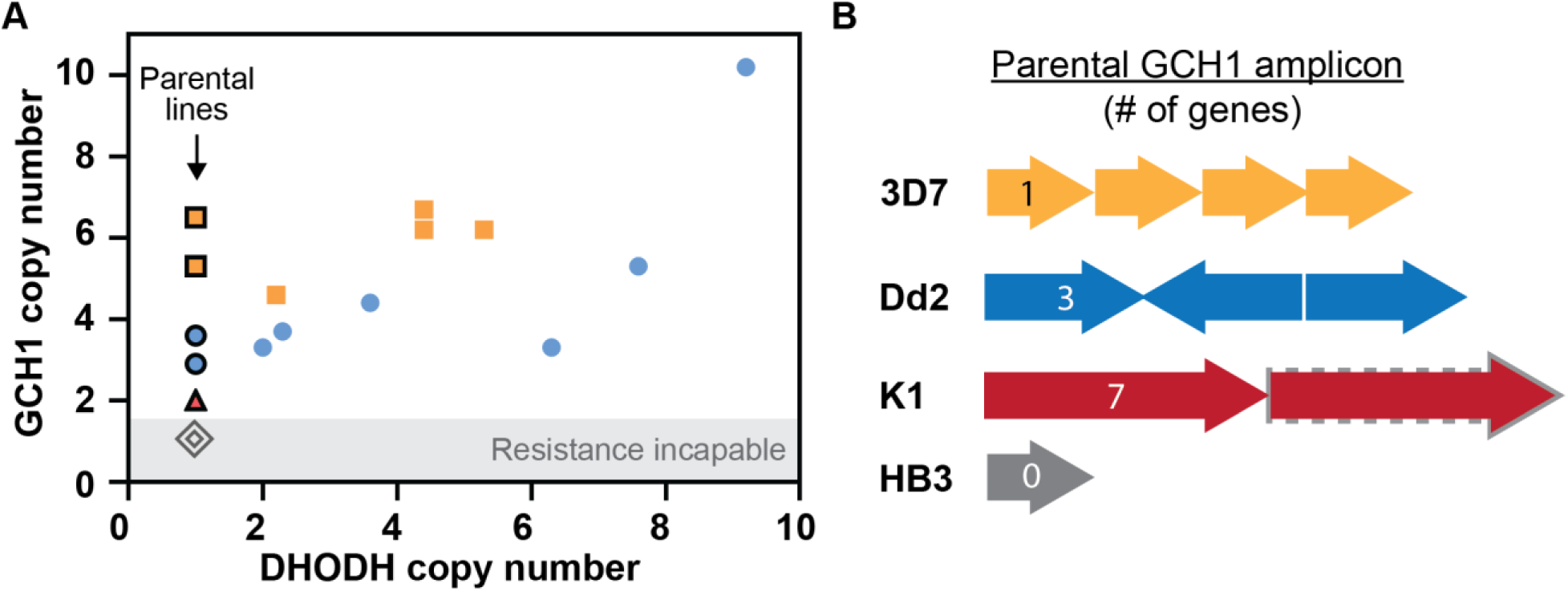
Preexisting *GCH1* amplicons confer resistance competence. (**A**) Relationship between *GCH1* and *DHODH* copy number in parental (black outline) and DHODH inhibitor-selected parasite lines as quantified using short read data (see **Table 2** for source data). Each data point represents a copy number from an individual parasite clone or line from a distinct selection (see **Tables S3** and **S4** for copy number assessment). *3D7*, yellow squares (2 selections); *Dd2*, blue circles (2 selections); *K1*, red triangle (1 selection, unknown *DHODH* copy number post-selection); *HB3*, grey diamonds (2 selections). (**B**) *GCH1* amplicon size and gene number vary between parasite lines used for DHODH inhibitor selections (see **Table 2** for selection details). *3D7*, 4 copy, single gene (PF3D7_1224000, 2kb, this study); *Dd2*, 3 copy, three genes (PF3D7_1223800-PF3D7_1224000, 5kb, this study); *K1*, unknown copy number (grey dash, anticipated 2 copy), 7 genes (PF3D7_1223500 - PF3D7_1224100, 19kb) (39); *HB3*, 1 copy of region (40).

**Table 2.** Parental line *GCH1* copy number determines resistance competence.

| Study<br>(author, year) | Selection<br>Compounds | Success of Resistance Selection |  |  |  | *DHODH*<br>amplicon |
| --- | --- | --- | --- | --- | --- | --- |
|  |  | *Dd2*<br>(*GCHI*=3) | *3D7*<br>(*GCHI*=4-6) | *Hb3*<br>(*GCHI*=1*) | Other<br>lines |  |
| Guler, 2013 (11) | DSM1 | Yes | N/A | No | N/A | Yes |
| Palmer, 2021 (37) | DSM705 | Yes | Yes | N/A | N/A | Yes |
| Phillips, 2015 (38) | DSM265 | Yes | N/A | No | K1-Yes** | Yes |
| Mandt, 2019 (18) | DSM265,<br>DSM267 | Yes | Yes | N/A | N/A | ^Yes |
| White, 2019 (41) | DSM265 | Yes | N/A | N/A | N/A | No |
| Ross, 2014 (42) | DSM74 | Yes | Yes | N/A | N/A | No |
| Lukens, 2014 (43) | Genz-666136,<br>DSM74 | N/A | Yes | N/A | N/A | No |

\*The *GCHI* copy number for the *HB3* parasite line is single copy based on whole genome sequencing (40). Prior studies detected two *GCHI* copies using quantitative PCR but did not detect an increase in expression from this region (8,15).
\*\*The *K1* parasite line has a *GCHI* amplicon that is ~19kb (961,455–980,630, (39)) but the exact copy number has not been reported. Phillips et al. reported that it was 100x more difficult to select for DSM265 resistance using *K1* than *Dd2* (MIR- 2 x 10<sup>8</sup> versus 2 x 10<sup>6</sup>, respectively, (38). Resistance mechanisms (i.e. mutation or amplification) were not evaluated for the resistant *K1* line in this study.
^Although both *Dd2* and *3D7* were used in selections, only two *3D7* clones acquired *DHODH* amplifications in this study (18) and were evaluated for *GCHI* copy number in our study (Tables S3 and S4).

## Discussion

Using both short-read sequencing as well as an accurate PCR-based method, ddPCR, we uncovered that *GCH1* copy number is positively associated with *DHODH* copy number in malaria parasites selected with DSM1 (**Figures 1B and 4**, **Table 1**). However, limitations in each of these methods contribute to inaccuracies in copy number results. Most importantly, both of these methods provide an average value for the parasite population and do not reflect variation from individual genomes. Long-read sequencing combined with a custom visualization tool (44) allowed us to directly quantify the copy number of the *GCH1* locus and assess the structure of all amplicons (**Figures 2** and **3**). In the context of this haploid, single cellular parasite, each read represents a distinct DNA strand from a single genome; this approach is an accurate way to visualize copy number and junction heterogeneity across a population (**Figure 2**, **Figure S2**). We showed that the detection of more *GCH1* amplicons on reads from DSM1 selected parasite lines (**Figure 3A**) was not caused by sample preparation (**Figure 3B**, **Table S2**) or natural variation during in vitro culture (**Table S2**). The visualization of tandem amplicons on single reads (**Figure 2A-C**) and the step-wise increase by 2 amplicon units (from 3 to 5 to 7 copies, **Figures 2D** and **S1**) provides direct evidence for *GCH1* amplicon expansion in the DSM1 resistance context.

The two-unit steps suggest that there is a secondary recombination site that leads to facile re-amplification of the region (**Figure 2D**). In the high coverage M1 sample, we identified long AT-rich sequences at the end of the amplicon unit (**Figure S4**) that may participate in further DNA breakage (AT dinucleotides at the *GCH1* end, **Figure S4C**) and recombination after the initial event (homopolymeric A/T tracks at the *YHM2* end, **Figure S4A**)(24). Multiple junctions at these positions provide evidence of the initial event that formed the 3 unit amplicon as well as a secondary event resulting in a two-unit step-wise expansion.

As recently recognized in organisms from bacteria to humans (34,35,45–47), long read sequencing is an exceptional tool to investigate genomic heterogeneity within populations. This method is not without its limitations; reads that end within the amplicon region lead to an underestimation in the total copy number; in our study, 65/98 (66%) of reads were non-spanning, which contributed to a similar overall mean copy number between parasite lines (**Figure 3A**). However, the visualization of spanning reads overcame this limitation, clearly showing the two-unit expansion of *GCH1* amplicons (**Figures 2A-C** and **S1**). The quantification of spanning reads also confirms that variation in *GCH1* amplicon numbers likely arises stochastically in the parasite population; while most spanning reads from the high coverage M1 line showed 5 amplicon copies (15/21 reads), we observed higher amplicon copies on a minority of reads (7 and 9 copies, 3/21 reads). In line with our previous model for malaria genome evolution (11,24), parasites with higher copy numbers are present within the population, poised to be selected under specific conditions.

The *GCH1* locus is a proven hotspot for genetic change (8,39). *GCH1* amplicons have previously been associated with antifolate resistance; an increase in flux through the folate biosynthesis pathway via *GCH1* alleviates fitness effects of mutations that confer pyrimethamine and sulfadoxine resistance (4,8,16,19,20). A similar contribution to resistance to DHODH inhibitors would not be surprising, given the close connection of the folate and pyrimidine biosynthesis pathways (**Figure 5**); they both contribute to nucleotide biosynthesis and converting dUMP to dTMP in pyrimidine biosynthesis requires reciprocal conversion of N^5^, N^10^-methylene-THF to DHF by the folate pathway. Strong evidence for this metabolic connection comes from the observation that parasite lines without *GCH1* amplicons are not able to develop resistance to DHODH inhibitors (**Table 2**) (11,38). Assessment of resistance competence, combined with our observations in the DSM1 context and folate-pyrimidine metabolic interplay, lead us to propose that increases in *GCH1* copy number are also beneficial for the fitness of parasites carrying *DHODH* amplicons. During resistance evolution, increased DHODH protein levels interact with more of the inhibitor to cause resistance. When the inhibitor is not present, higher DHODH levels increase flux through pyrimidine biosynthesis, which in turn requires more THF for the completion of the pathway (**Figure 5**).

**Fig 5.**
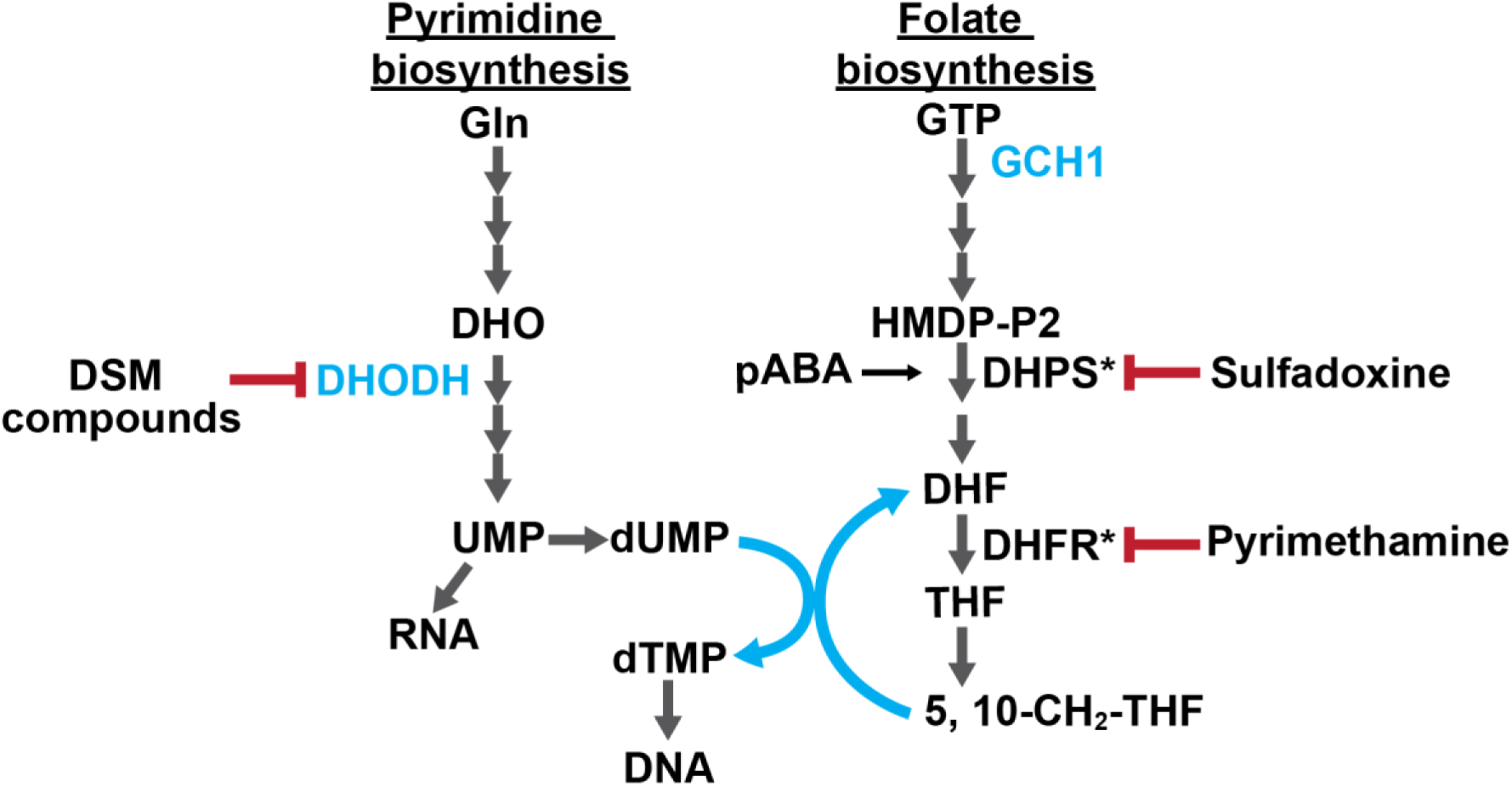
The connection between pyrimidine and folate biosynthesis pathways. Enzymes with gene copy number variations are indicated in blue text (DHODH: dihydroorotate dehydrogenase; GCH1: GTP cyclohydrolase 1) and connections between the two pathways in blue arrows. Gln: Glutamine; DHO: Dihydroorotate; UMP: Uridine monophosphate; dUMP: Deoxyuridine monophosphate; dTMP: deoxythymidine monophosphate; GTP: Guanosine-5’-triphosphate; DHPS: Dihydropteroate synthase; DHFR: Dihydrofolate reductase; DHF: Dihydrofolate; THF: Tetrahydrofolate; HMDP-P2: 6-hydroxymethyl-7, 8-dihydropterin diphosphate; pABA: para-amino-benzoic acid; 5, 10-CH2-THF: 5,10-Methylenetetrahydrofolate. \**Dd2* carries 3 mutations in both DHPS and DHFR (6 total) and HB3 and 3D7 have a single mutation each in DHFR and DHPS, respectively (15,48).

There are a few factors that may impact parasite reliance on folate biosynthesis including parasite genetic background and the presence of folate precursors in the host environment. Antifolate resistant parasites from different regions of the world that harbor specific mutations rely more heavily on increased GCH1 flux to alleviate fitness effects (i.e. DHFR and DHPS mutants, (4,8,15,19), **Figure 5**). While *Dd2* and *3D7* have distinct origins and mutational patterns (49,50) (**Figure 5**), the fact that they both maintain the ability to develop DHODH inhibitor resistance excludes the possibility that DHFR and DHPS mutations matter in this context. Additionally, levels of folate precursors, such as para-aminobenzoic acid (pABA), vary widely in host versus in vitro environments (51–55) and exert different selective pressures on parasites during drug selection (56). Higher pABA levels in human serum, as were used during DSM1 selections (11), may drive the accumulation of higher *GCH1* copy numbers and hold extra benefits by bolstering both folate and pyrimidine biosynthesis (**Figure 5**). Our observation of some level of variation in *GCH1* amplicon number in standard parasite lines (**Table S2**) suggests that differing in vitro media formulations may drive copy number changes at this locus.

Importantly, DHODH inhibitors have entered trials for the treatment of clinical malaria (18) and resistance-conferring *DHODH* amplicons can evolve in vivo (57). Meanwhile, antifolate resistance and *GCH1* amplifications are widespread in clinical *P. falciparum* isolates, especially in Southeast Asia (4,7,8,39) and now Africa (16,58). Additionally, a zoonotic malaria species, *P. knowlesi,* also acquires extra *GCH1* copies after adaptation to human blood (59). The potential use of DHODH inhibitors on clinical parasites carrying *GCH1* amplicons necessitates further evaluation of the interplay between these two important metabolic pathways.

## Materials and Methods

### DSM1 and parasite clones

DSM1 inhibits dihydroorotate dehydrogenase enzyme of *Plasmodium* pyrimidine biosynthesis (60). DSM1-resistant parasites were selected during a previous study and cloned by limiting dilution at the time of isolation (11) (**Figure 1A**). The naming scheme representing low (L), moderate (M), and high (H) levels of DSM1 resistance follows that used previously by our group (36).

### Parasite culture

We thawed erythrocytic stages of *P. falciparum* (*Dd2*, MRA-150; *3D7*, MRA-102, Malaria Research and Reference Reagent Resource Center, BEI Resources and DSM1 selected clones as highlighted in **Figure 1A**, a generous gift from Pradipsinh Rathod, University of Washington) from frozen stocks and maintained them as previously described (61). Briefly, we grew parasites at 37°C in vitro at 3% hematocrit (serotype A positive human erythrocytes, Valley Biomedical, VA or BioIVT, NY) in RPMI 1640 medium (Invitrogen, USA) containing 24 mM NaHCO3 and 25 mM HEPES, and supplemented with 20% human type A positive heat inactivated plasma (Valley Biomedical, VA or BioIVT, NY) in sterile, sealed flasks flushed with 5% O_2_, 5% CO_2_, and 90% N_2_ (11). We maintained the cultures with media changes every other day and sub-cultured them as necessary to keep parasitemia below 5%. We determined all parasitemia measurements and staging using SYBR green-based flow cytometry (62). We routinely tested cultures using the LookOut Mycoplasma PCR Detection Kit (Sigma-Aldrich, USA) to confirm negative Mycoplasma status.

### Short read sequencing analysis and CNV detection

We analyzed amplifications in Illumina short read datasets of *P. falciparum* parasites selected by different DSM antimalarial drugs (DSM1, DSM265, DSM267, and DSM705, **Table S3**) (11,18,37). We first processed and mapped the reads to the reference genome as previously described (24,63). Briefly, we trimmed Illumina adapters from reads with the BBDuk tool in BBMap (version 38.57) (64). We aligned each fastq file to the *3D7* reference genome with Speedseq (version 0.1.0) through BWA-MEM alignment (65). We discarded the reads with low-mapping quality scores (below 10) and removed duplicated reads using Samtools (version 1.10) (66). We analyzed split and discordant reads from the mapped reads using LUMPY in Speedseq to determine the location and length of the previously reported *GCH1*, *DHODH*, and multidrug resistance protein 1 (*MDR1*) amplicons (**Table S4**) (67). For read-depth analysis, we further filtered the mapped reads using a mapping quality score of 30. To determine the copy number of the *GCH1*, *DHODH*, and *MDR1* amplicons, we used CNVnator (version 0.4.1) with a bin size of 100 bp; the optimal bin size was chosen to detect GCH1 amplicons in all analyzed samples (68).

### Droplet Digital PCR

Prior to Droplet Digital (dd) PCR, we digested DNA with restriction enzyme RsaI (Cut Site: GT/AC) following the manufacturer’s instructions (New England Biolabs, Ipswich, MA, USA) in 37°C incubation for one hour. We selected the restriction enzyme RsaI to cut outside of the ddPCR amplified regions of desired genes and separate amplicon copies to be distributed into droplets. We diluted the digested DNA for ddPCR reactions. We performed ddPCR using ddPCR Supermix for Probes (no dUTP, Bio-Rad Laboratories, California, USA) with DNA input 0.1 ng (in duplicate per sample), 0.025 ng (in duplicate per sample) as previously described (36). The primers and probes used in reactions are included in **Table S5**. The PCR protocol for probe-based assay was 95°C for 10 min, followed by 40 rounds of 95°C for 30 sec and 60°C for 1 min. *Seryl-tRNA synthetase* and *calcium-transporting ATPase* (*ATP6*) served as single copy reference genes on chromosome 7 and chromosome 1 respectively; *dihydroorotate dehydrogenase* (*DHODH*) and *GTP cyclohydrolase 1* (*GCH1*) are multi-copy genes (**Table S5**). We performed droplet generation and fluorescence readings per the manufacturer’s instructions. For each reaction, we required a minimum number of 10,000 droplets to proceed with analysis. We calculated the ratio of positive droplets containing a single- (*ATP6*) or multi-copy gene (*GCH1*, *DHODH*) versus a single-copy gene (*seryl-tRNA synthetase*) using the Quantasoft analysis software (QuantaSoft Version 1.7, BioRad Laboratories) and averaged between independent replicates.

### DNA extraction for long read sequencing

We lysed asynchronous *P. falciparum*-infected erythrocytes with 0.15% saponin (Sigma-Aldrich, USA) for 5 min at room temperature and washed them three times with 1× PBS (diluted from 10× PBS Liquid Concentrate, Gibco, USA). We then lysed parasites with 0.1% Sarkosyl Solution (Bioworld, bioPLUS, USA) in the presence of 1 mg/ml proteinase K (from *Tritirachium album*, Sigma-Aldrich, USA) overnight at 37°C. We first extracted nucleic acids with phenol/chloroform/isoamyl alcohol (25:24:1) pH 8.0 (Sigma-Aldrich, USA) three times using 1.5ml light Phase lock Gels (5Prime, USA), then further extracted nucleic acids with chloroform twice using 1.5ml light Phase lock Gels (5Prime, USA). Lastly, we precipitated the DNA with ethanol using the standard Maniatis method (69). To obtain high molecular weight genomic DNA, we avoided any pipetting during the extraction, transferred solutions by direct pouring from one tube to another, and mixed solutions by gently inverting the tubes.

### Oxford Nanopore long read sequencing and analysis

For WT1 and H2 samples, we subjected ~1μg of high molecular weight genomic DNA to library preparation for Oxford Nanopore sequencing following the Nanopore Native barcoding genomic DNA protocol (version: NBE_9065_v109_revAB_14Aug2019) with 1x Ligation Sequencing kit (SQK-LSK109, Oxford Nanopore Technologies, Oxford, UK). We performed DNA repair and end preparation using NEBNext FFPE DNA Repair Mix (New England Biolabs, Ipswich, MA, USA) and NEBNext End Repair/dA-Tailing Module (New England Biolabs, Ipswich, MA, USA). We cleaned the A-tailed fragments using 0.9X AMPure XP beads (Beckman Coulter, High Wycombe, UK). We then ligated barcodes to the end-prepped DNA using the Native Barcoding Expansion 1–12 kit (EXP-NBD104, Oxford Nanopore Technologies, Oxford, UK) and Blunt/TA Ligase Master Mix (New England Biolabs, Ipswich, MA, USA). We cleaned the barcoded samples using 0.9X AMPure XP beads. Then we pooled barcoded samples in equimolar ratios and subjected them to an adaptor ligation step, using the Adapter Mix II from the Native Barcoding Expansion 1–12 kit and NEBNext Quick Ligation Reaction Buffer (New England Biolabs, Ipswich, MA, USA) as well as Quick T4 DNA Ligase (New England Biolabs, Ipswich, MA, USA). After adaptor ligation, we cleaned the WT1 and H2 libraries using AMPure XP beads.

For M1 samples, we isolated high molecular weight genomic DNA from 10 flasks of parasites (1-4% parasitemia, ~30-40% late stage parasites) and prepared libraries as above except that we used the cleanup and precipitation reagents provided in the Ultra-long DNA Sequencing kit (Oxford Nanopore Technologies, Oxford, UK, V14). We quantified the adapter-ligated and barcoded DNA using a Qubit fluorimeter (Qubit 1X dsDNA High Sensitivity Assay Kit, Life Technologies, Carlsbad, CA).

We sequenced the WT1 and initial H2 libraries using the R9.4.1 flow cell (FLO-MIN106D), another H2 library using the R10 flow cell (FLO-MIN111), and M1 libraries using the R10.4.1 flow cell (FLO-MIN114) on MinION (Oxford Nanopore Technologies, Oxford, UK). For WT and H2 samples, we ran flow cells for 48 hours (controlled and monitored using the MinKNOW software 3.6.5). For the M1 samples, we ran one-third of the library sequentially on the flow cell for 24 hours each (up to 72 hours in total).

For base calling and demultiplexing of the Nanopore sequencing reads, we used Guppy (version 3.4.5+fb1fbfb) with the parameter settings “-c dna_r9.4.1_450bps_hac.cfg --barcode_kits “EXP-NBD104” -x auto” for samples sequenced with R9.4.1 flow cell and “-c dna_r10_450bps_hac.cfg -x auto” for the sample sequenced with R10 flow cell. We checked the read length and read quality using “Nanoplot” (version 1.0.0) (see **Table S1**). We trimmed the adapters with “qcat” (version 1.1.0) (70) and filtered the reads with a cutoff “length ≥ 500 and Phred value ≥ 10” using the program “filtlong version 0.2.0” (https://github.com/rrwick/Filtlong). To estimate the coverage of sequencing reads in each sample, we aligned the filtered reads to *3D7* reference genome (PlasmoDB) using “minimap2” (version 2.17) (71). “QualiMap” (version 2.2.1) (72) was used to calculate the coverage of the aligned reads (see **Table S1**).

For junction analysis, we aligned Q10-filtered M1 reads (.bam) to the *3D7* reference genome (PlasmdDB) and visualized read sequence at junctions of the GCH1 amplicon using Integrated Genome Viewer (73,74).

### Direct visualization of long reads

To visualize structural variants in the parasite genome, we used a custom R Shiny script to plot the arrangement of reference gene segments on individual Nanopore reads (44). Briefly, we defined a target region in the *3D7* reference genome (chromosome 12: 932916bp – 999275bp) that contained 3 genes in the *GCH1* amplicon and 11 flanking genes. We extracted the reference sequences of these genes and subsequent intergenic regions, then split these sequences into fragments of 500-1000 base pairs. We compared these fragments to individual Nanopore reads (>10kb) using BLAST (75). We used the BLAST output as input for a custom R script, which drew blocks representing homology between the defined genes (y-axis) and each individual read (x-axis). If genes on the read were arranged in the same order as the reference genome then the blocks would create a diagonal line on the plot. If there was a CNV that altered the gene order or direction then the diagonal line is disrupted. We allowed the percent identity required to draw a homologous rectangle to vary between reads, which varied in quality, using a slider in the Shiny application. To filter out long reads with potentially spurious hits to gene fragments, we also used BLAST to compare Nanopore reads to the reference genome and removed reads with <90% identity to chromosome 12. To compare the mean copy number of GCH1 amplicon and read lengths between WT1, M1, and H2 reads, we performed a one-way ANOVA in Graphpad Prism with an alpha value of 0.05. If significant, differences were further explored using Tukey’s multiple comparisons testing.

## Acknowledgments

Our thanks to Michelle Warthan, Noah Brown, and Ali Guler (University of Virginia) for experimental, analysis, and statistical support, Martin Wu and Mercedes Campos-Lopez (University of Virginia) for Oxford Nanopore sequencing support, and the laboratory of Dr. William Petri Jr (University of Virginia) for their helpful discussions and insight. We are grateful to David Fidock (Columbia University) and Dyann F. Wirth (Harvard University) for sharing sequencing data and information on DSM-selected cell lines.

## Supporting Information Captions

Figure S1: Spanning reads carry odd copy numbers of *GCH1* amplicons

Figure S2: Long-read visualization of *GCH1* amplicon from *3D7* parasite line

Figure S3: Parasites lines with *DHODH* amplicons are selected from parent lines with *GCH1* amplicons

Figure S4: *GCH1* amplicon junctions show signs of primary and secondary expansion events

Supplemental Methods: Procedures used for the preparation of long read data by Ellen Yeh’s group, Stanford University, as presented in Table S2, including in vitro cultures, mutagenesis, DNA extraction, and sequencing.

Table S1: Oxford Nanopore sequencing datasets new to this manuscript

Table S2: Quantification of long reads covering the *GCH1* amplicon displays greater variability in amplicon number from DSM1 resistant parasites

Table S3: *DHODH* and *GCH1* copy number in parasites resistant to DHODH inhibitors

Table S4: Detection of known amplifications by Illumina short reads from parasites resistant to DHODH inhibitors

Table S5: ddPCR primer and probe sequences

