## Supplementary material for "Direct long read visualization reveals metabolic interplay between two antimalarial drug targets": Figure S

**Supporting Information**

Supplementary Figures

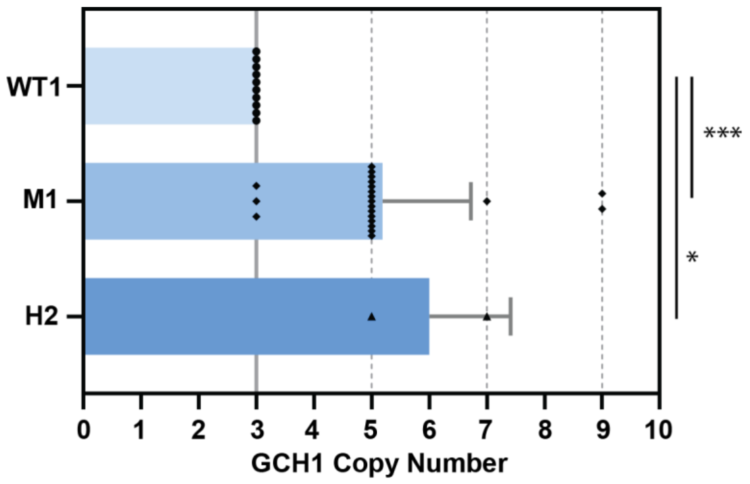


**Figure S1. Spanning reads carry odd copy numbers of *GCH1* amplicons.** Only spanning reads, or those that cover both ends of the amplicon as well as some up- and down-stream genes, were included in this analysis. Overall, 33 of 98 total reads were characterized as spanning reads (WT1, 10/19 reads; M1, 21/62 reads; H2, 2/18 reads). ***, $p<0.0005$. *, $p <0.05$ (Tukey's multiple comparisons test)

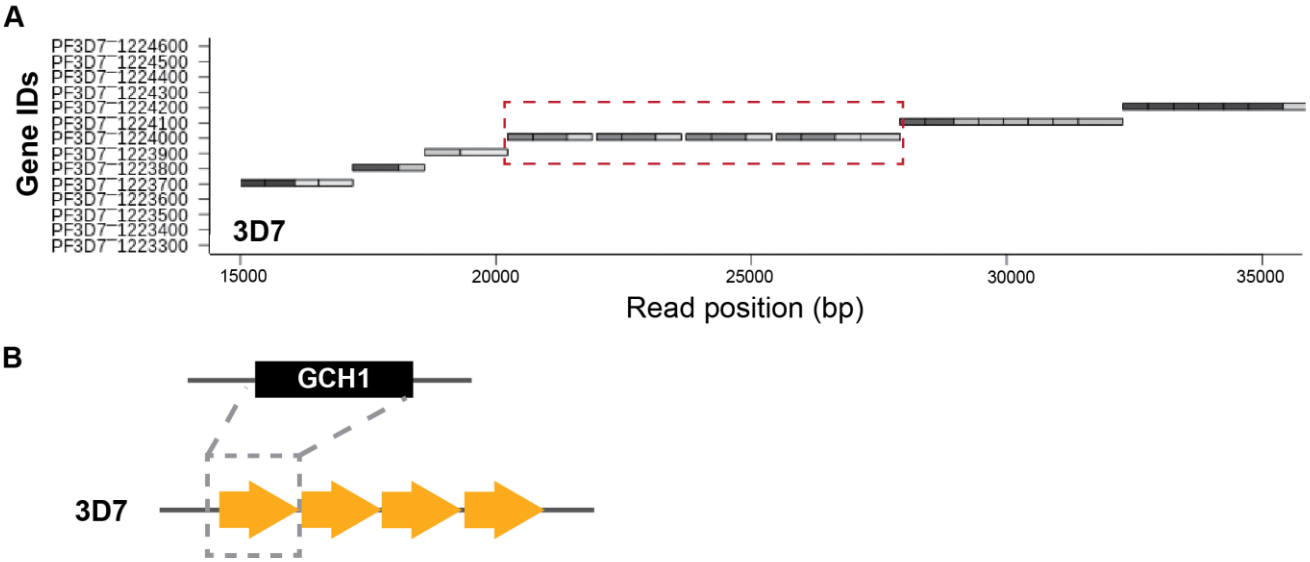


**Figure S2. Long-read visualization of *GCH1* amplicon from *3D7* parasite line. (A)** Representative image from the Shiny application of the *GCH1* amplicon from the *3D7* genome (see sequencing results in **Tables S1 and S2**). Red dashed square: *GCH1* amplicon covering 1 gene. Each gene sequence from the *3D7* reference genome (no *GCH1* amplicon represented) was split into <=500 bp fragments and blasted against individual Nanopore reads (dark gray: genic regions, light grey: intergenic regions, more details in *Materials and Methods*). **(B)** Orientation of *GCH1* amplicon from the *3D7* parasite line. The one gene within the *GCH1* amplicon unit is PF3D7_1224000 (*GTP cyclohydrolase I, GCH1*).

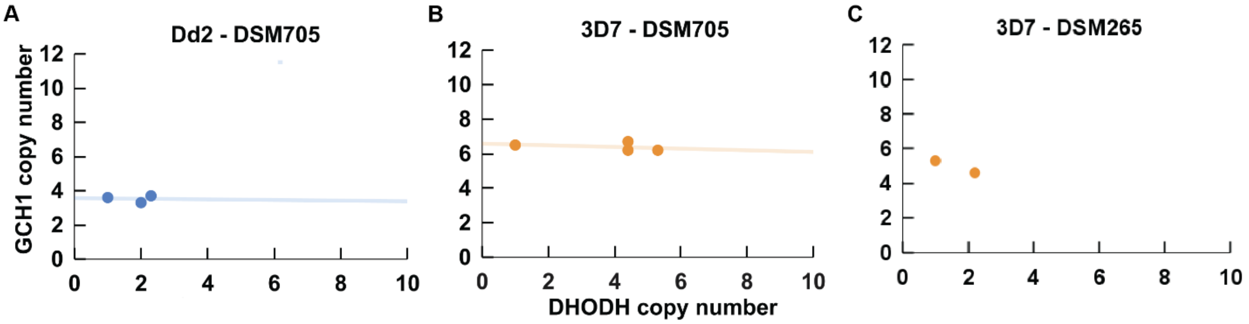


**Figure S3. Parasites lines with *DHODH* amplicons are selected from parent lines with *GCH1* amplicons**. Trend lines show the relationship between *GCH1* and *DHODH* copy numbers in available lines. **(A)** Relationship between *GCH1* and *DHODH* copy number in *Dd2* parent line and DSM705-selected lines as quantified using short read data (Palmer et al., 2021). **(B)** Relationship between *GCH1* and *DHODH* copy number in *3D7* parent line and DSM705-selected lines as quantified using short read data (Palmer et al., 2021). **(C)** Relationship between *GCH1* and *DHODH* copy number in *3D7* parent line and DSM265-selected lines as quantified using short read data (Mandt et al., 2019). Correlation coefficients are not calculated due to small sample sizes and dependence among selected lines.

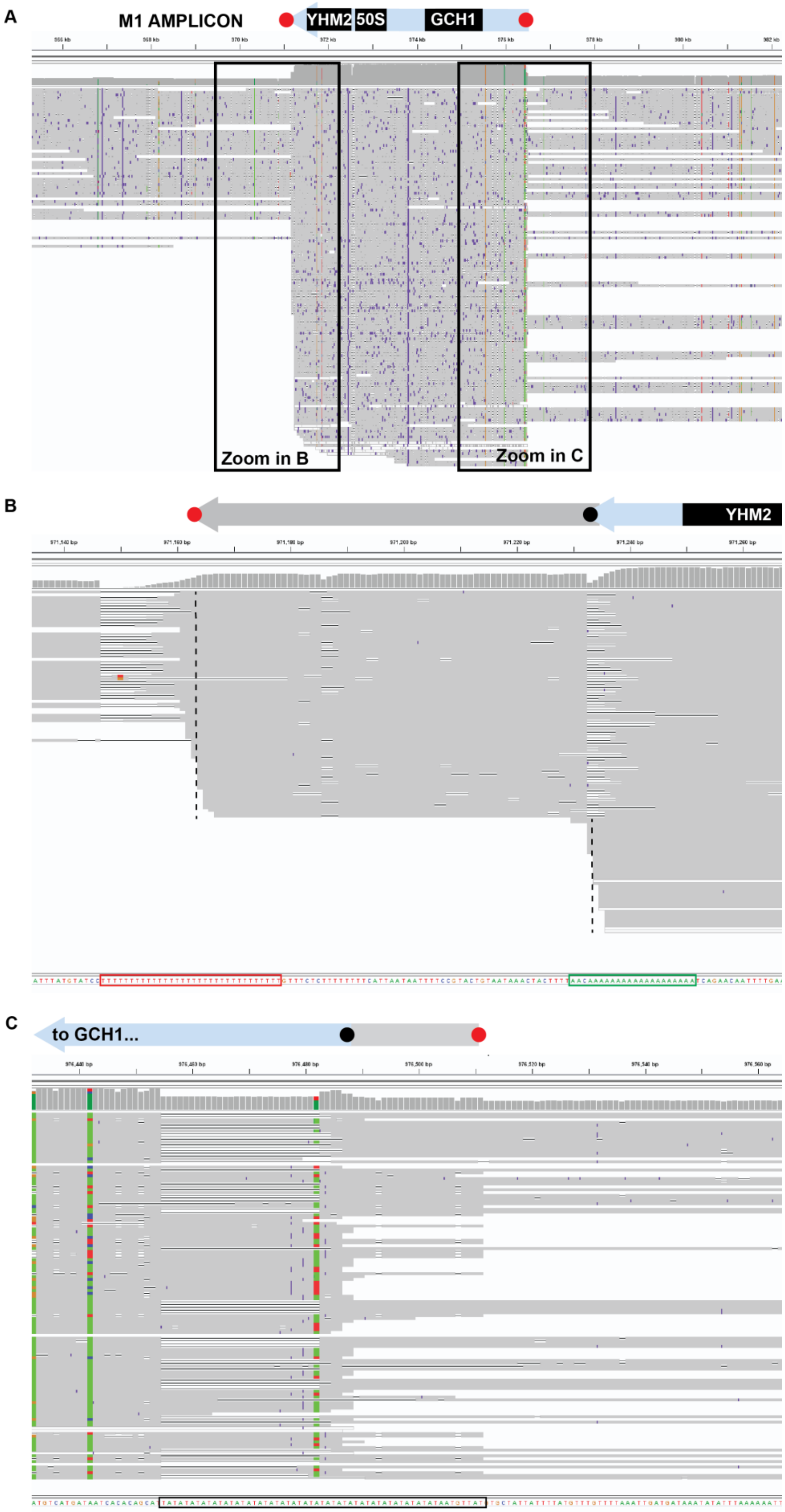
A
M1 AMPLICON
YHM2
50S
GCH1
Zoom in B
Zoom in C
B
YHM2
C
to GCH1...

**Figure S4.** ***GCH1*** **amplicon junctions show signs of primary and secondary expansion events.** M1 reads were aligned to the *3D7* reference using Integrated Genome Viewer (amplicon orientation is consistent with the reference genome but opposite from Figure 2). For simplicity of viewing, the *3D7* reference genome was used for alignment because it does not reflect the *GCH1* amplicon (unlike the parental *Dd2* genome, which reflects 2 inverted *GCH1* amplicons). The amplicon location and direction (blue arrows) are depicted at the top of each panel with approximate locations of genes (black boxes) and primary and secondary amplicon junctions (red and black circles). Grey reads, regions that are homologous to consensus; purple regions, insertion-deletions relative to the reference genome; colored bases, conflicting bases relative to the reference genome. The consensus sequence is shown at bottom of each panel. (**A**) Entire amplicon region. Reads that end at the junction represent those that span the amplicon junction. Reads that extend beyond this region represent those that extend into neighboring genes. (**B** and **C**) Zoom of upstream and downstream ends of the amplicon (relative to *3D7* reference). Positions where reads end indicate amplicon junction sites (dashed line). Green/red outline, stretches of homopolymeric A or T tracks in the consensus sequence; black outline, AT dinucleotide track. The presence of secondary junctions (black circles), or those within the primary junctions (red circles), indicate additional breakage and recombination sites following the initial event.

### Supplementary Methods

Procedures used for the preparation of long read data by Elen Yeh's group, Stanford University, as presented in **Table S2**

#### In vitro cultures

*Plasmodium falciparum* parasites W2 (MRA-157), Dd2 (MRA-150), and 3D7 (MRA-102) were obtained from MR4. Parasites were grown in human erythrocytes (Stanford Blood Center) at 2% hematocrit in RPMI 1640 medium (Gibco) supplemented with 0.25% Albumax II (Gibco), 2 g/liter sodium bicarbonate (Fisher), 0.1 mM hypoxanthine (Sigma), 25 mM HEPES, pH 7.4 (Sigma), and 50 µg/liter gentamicin (Gold Biotechnology) at 37°C, 5% O2, and 5% CO2.

#### Mutagenesis

Synchronized 3D7 late-stage parasites were incubated in complete medium with 6.17uM N-ethyl-N-nitrosourea (Sigma) for 2hrs. Parasites were washed 3 times with complete media and separated into multiple flasks at approximately 10% parasitemia, 2% HTC. Cultures were maintained for a month before gDNA was extracted.

#### DNA extraction

Parasites from the original clone W2 and Dd2 (W2_PC, Dd2_PC), selected resistant clones via pathogens box from W2 and Dd2 (W2_C1, Dd2_674140), and mutagenized 3D7 (Plasmodium_3D7_6uM) were isolated from erythrocytes by addition of 0.1% saponin (Sigma) maintained on ice for 5 min. Lysed parasite pellets were washed three times with ice cold phosphate-buffered saline to remove extracellular DNA. DNA was extracted using either the Circulomics Blood and Tissue kit (Pacific BioSciences) following manufacturer's instructions or by phenol-chloroform-isoamyl (25:24:1). In brief the phenol-chloroform-isoamyl extraction were preceded by overnight lysis of parasites in phosphate buffered saline with 0.1% L—loril sarkosil (Sigma) and 200 mg/ml proteinase K. Nucleic acids were then extracted with phenol-chloroform-isoamyl (25:24:1) followed by RNA digestion with 100 mg/ml RNase A for 1 hour at 37C. gDNA was extracted twice more with phenol-chloroform-isoamyl alchol and once with 100% chloroform. DNA was precipitated with a 2.5X volume of 100% ethanol followed by two washes with 70% ethanol. gDNA was stored at 4C prior to sequencing.

#### Sequencing

We subjected 1ug if HMW DNA for each sample to library preparation using Oxford Nanopore sequencing following the genomic DNA by ligation protocol (Version GDE_9063_v109_revS_14Aug2019 for SQK-LSK109 libraries; GDE_9108_v110_revC_10Nov2020 for SQK-LSK110 libraries) with the genomic DNA by ligation kits (SQK-LSK109; SQK-LSK110) with extended times for end repair and ligation. We performed DNA repair and end preparation using NEBNext FFPE DNA Repair Mix (New England Biolabs, Ipswich, MA, USA) and NEBNext End Repair/dA-Tailing Module (New England Biolabs, Ipswich, MA, USA) extending times for 5 to 30 minutes for the end repair (for both the 20C and 65C incubation). We cleaned the A-tailed fragments using 0.9X AMPure XP beads (Beckman Coulter, High Wycombe, UK). We then ligated adapters to the end-prepped DNA using the NEBNext quick ligation modules (New England Biolabs, Ipswich, MA, USA) with a ligation extended from 10 to 60 minutes. After adaptor ligation, we cleaned the library using AMPure XP beads and washed the library using the long fragment buffer (LFB; Oxford Nanopore Technologies, Oxford, UK). We quantified the adapter ligated DNA using a Qubit fluorimeter (Qubit 1X dsDNA High Sensitivity Assay Kit, Life Technologies, Carlsbad, CA). We sequenced all libraries using the R10 flow cell (FLO-MIN111) on MinION for 12-24 hours. Exact sequencing time is dependent on flow cell health and data from previously loaded library.
